## Supplementary material for "Characterizing collaborative transcription regulation with a graph-based deep learning approach"

| Type | Notation | Explanation |
|---|---|---|
| Subscripts & superscripts | $i$ | Index of a segment of DNA sequence, $i = 1, \ldots, N$ |
| | $l$ | Index of a chromatin feature, $l = 1, \ldots, L$ |
| | $c$ | Spatial relationship from a contact map |
| | $s$ | Sequential relationship from DNA sequences |
| Operators | $\|\|$ | Concatenation |
| | $T$ | Transpose |
| | $\cup$ | Union |
| Symbols | $\mathcal{V}$ | Set of nodes |
| | $\mathcal{E}$ | Set of edges |
| | $\mathcal{A}$ | Weighted adjacency matrix |
| | $\mathcal{G}$ | Graph: $\mathcal{G} = (\mathcal{V}, \mathcal{E}, \mathcal{A})$ |
| | $R$ | Set of real numbers |
| | $\mathcal{N}_s^{(i)}$ | Sequential neighbor set of sequence $i$ |
| | $\mathcal{N}_c^{(i)}$ | Spatial neighbor set of sequence $i$ |
| | $\mathcal{N}^{(i)}$ | Neighbor set of sequence $i$: $\mathcal{N}^{(i)} = \mathcal{N}_s^{(i)} \cup \mathcal{N}_c^{(i)}$ |
| Neural network layers & functions | $f$ | Sequence layers |
| | $g_c, g_s$ | Graph layers |
| | $p$ | Prediction layer |
| | SAMPLE | Functions of sampling sequences from the neighborhood |
| | STACK | Functions of stacking multiple vectors to a matrix |
| Random variables | $x^{(i)}$ | One-hot representation of 1000-bp DNA sequence |
| | $\hat{y}^{(i)}$ | Predicted chromatin feature vector of sequence $i$ |
| | $\hat{y}_l^{(i)}$ | Predicted chromatin feature $l$ on sequence $i$ |
| | $\phi^{(i)}$ | Hidden representation vector of sequence $i$ extracted by sequence layers $f$ |
| | $\Xi_s^{(i)}$ | The feature matrix of sequence $i$ as an input to graph layers $g_s$ |
| | $\Xi_c^{(i)}$ | The feature matrix of sequence $i$ as an input to graph layers $g_c$ |
| | $h_s^{(i)}$ | Updated hidden representation of sequence $i$ as an output from $g_s$ |
| | $h_c^{(i)}$ | Updated hidden representation of sequence $i$ as an output from $g_c$ |
| | $P_c^{(i)}$ | Spatial sampling matrix |
| | $P_s^{(i)}$ | Sequential sampling matrix |
| | $[S_c^{(i)}]_l$ | Attribution scores of spatial sampling matrix $P_c^{(i)}$ for chromatin feature $l$ |
| | $[S_s^{(i)}]_l$ | Attribution scores of sequential sampling matrix $P_s^{(i)}$ for chromatin feature $l$ |
| | $[V_c^{(i)}]_l$ | Compressed interaction importance vector from $[S_c^{(i)}]_l$ for chromatin feature $l$ |
| | $[V_s^{(i)}]_l$ | Compressed interaction importance vector from $[S_s^{(i)}]_l$ for chromatin feature $l$ |
| | $M_l$ | Interaction importance matrix for chromatin feature $l$ |
| | $[S^{(j)}]_l^{(i)}$ | Attribution scores of sequence $j$ for chromatin feature $l$ |
| Parameters to be prespecified | $k_c$ | Number of sampled spatial neighbors |
| | $k_s$ | Number of sampled sequential neighbors |
| | $L$ | Number of chromatin features |
| | $K$ | Length of hidden representation $\phi^{(i)}$ |
| | $N$ | Number of 1000-bp DNA sequence |

Table S1: Notations used in our work

| | Mean AUROC | Mean AUPR |
|---|---|---|
| DeepSEA | 0.881 | 0.312 |
| DanQ | 0.881 | 0.316 |
| DeepCNN | 0.885 | 0.318 |
| ECHO (built on DeepSEA) | 0.918 | 0.373 |
| ECHO (built on DanQ) | 0.919 | **0.386** |
| ECHO (built on DeepCNN) | **0.921** | 0.378 |

Table S2: Comparing the mean AUROC and AUPR scores of ECHO with the baselines. The first three models are the baselines and the last threes are our proposed methods with the corresponding baseline model as the pre-trained sequence layers.

| | Mean AUROC | Mean AUPR | Mean Recall at 50% FDR |
|---|---|---|---|
| DeepCNN | 0.900 | 0.377 | 0.330 |
| ChromeGCN | 0.916 | 0.406 | 0.372 |
| ECHO | **0.924** | **0.429** | **0.399** |

Table S3: Comparing the performance of ECHO with ChromeGCN for 103 chromatin features on GM12878 cell line.

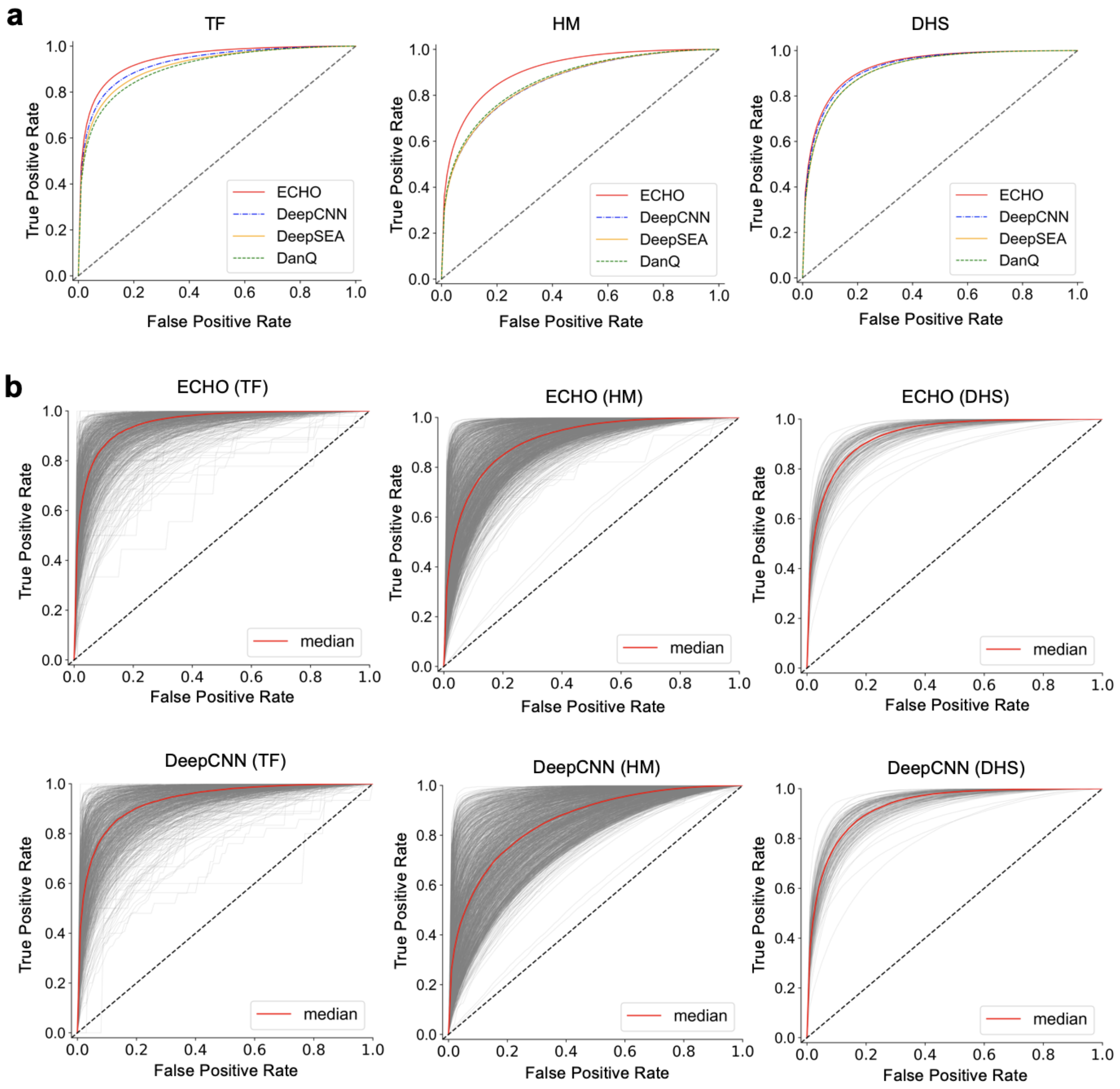


Figure S1: Comparing the prediction performance between ECHO and baselines.
(**a**) The mean ROC curves from ECHO and three baseline models for three types of chromatin features, including TF, HM and DHS. ECHO achieves higher mean AUROC scores than the baselines, especially on TF and HM. (**b**) The ROC curves for each chromatin feature from ECHO and DeepCNN models. The red lines denote the median ROC curves.

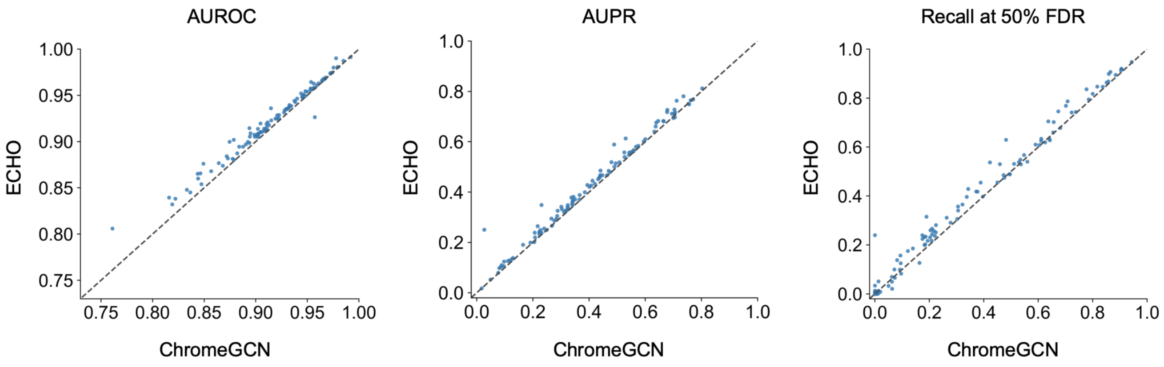


Figure S2: Scatter plots to compare the model performances of ECHO with ChromeGCN on GM12878 cell line.

CNN block (kernels, kernel size, stride, padding):

- Convolutional layer (kernels, kernel size, stride, padding)
- Batch normalization
- ReLU
- Convolutional layer (kernels, kernel size, stride, padding)
- Batch normalization
- ReLU

Graph layers $g_c$ and $g_s$:

1. CNN block (128, 20, 1, 0)
2. Max-pooling (size: 5)
3. CNN block (240, 10, 1, 0)
4. Max-pooling (size: 5)
5. Dropout (rate: 0.2)
6. CNN block (320, 10, 1, 0)
7. Max-pooling (size: 5)
8. Dropout (rate: 0.2)
9. CNN block (1300, 10, 1, 0)
10. Dropout (rate: 0.3)
11. Global average pooling

Prediction layer: one fully connected layer

Figure S3: Details of the graph layers and the prediction layer in ECHO.
The architectures of graph layers are varied considering the number of chromatin features, the input sequence size, and whether sequential neighbors are sampled. The model architecture reported here is for predicting 2,583 chromatin features with 50 spatial neighbors and 10 sequential neighbors per input sequence.

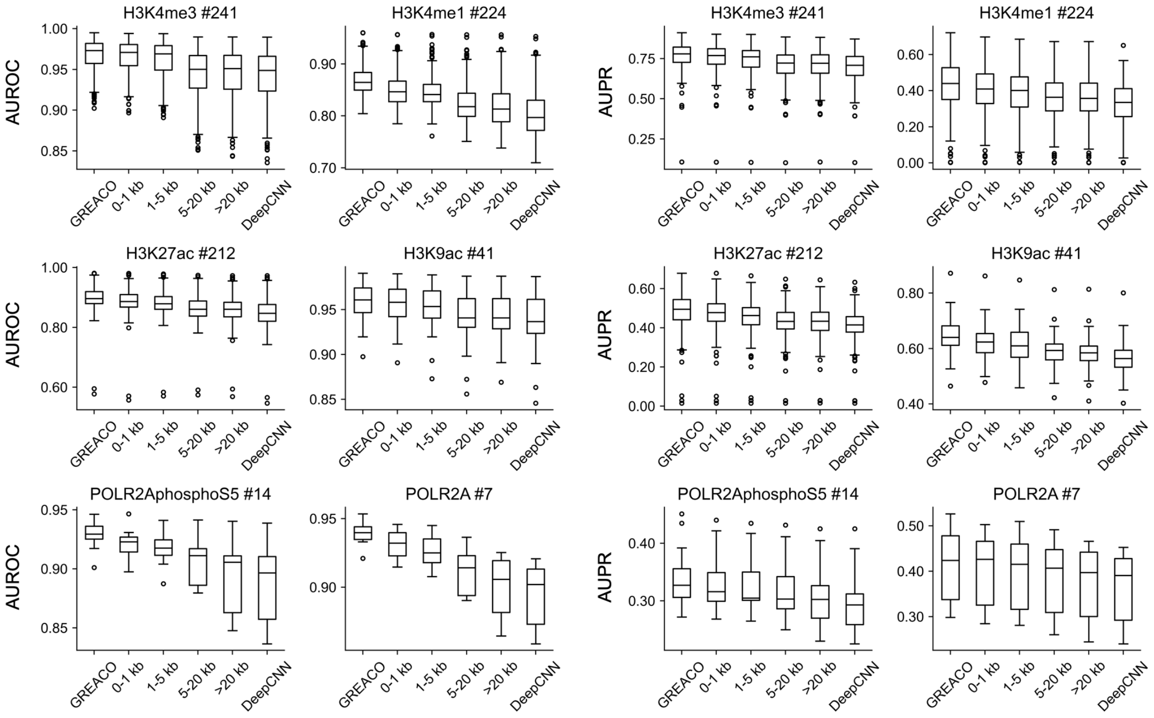


Figure S4: The effects of Micro-C contact distances on predicting chromatin features features mostly binding to promoters and enhancers and related with gene activation.

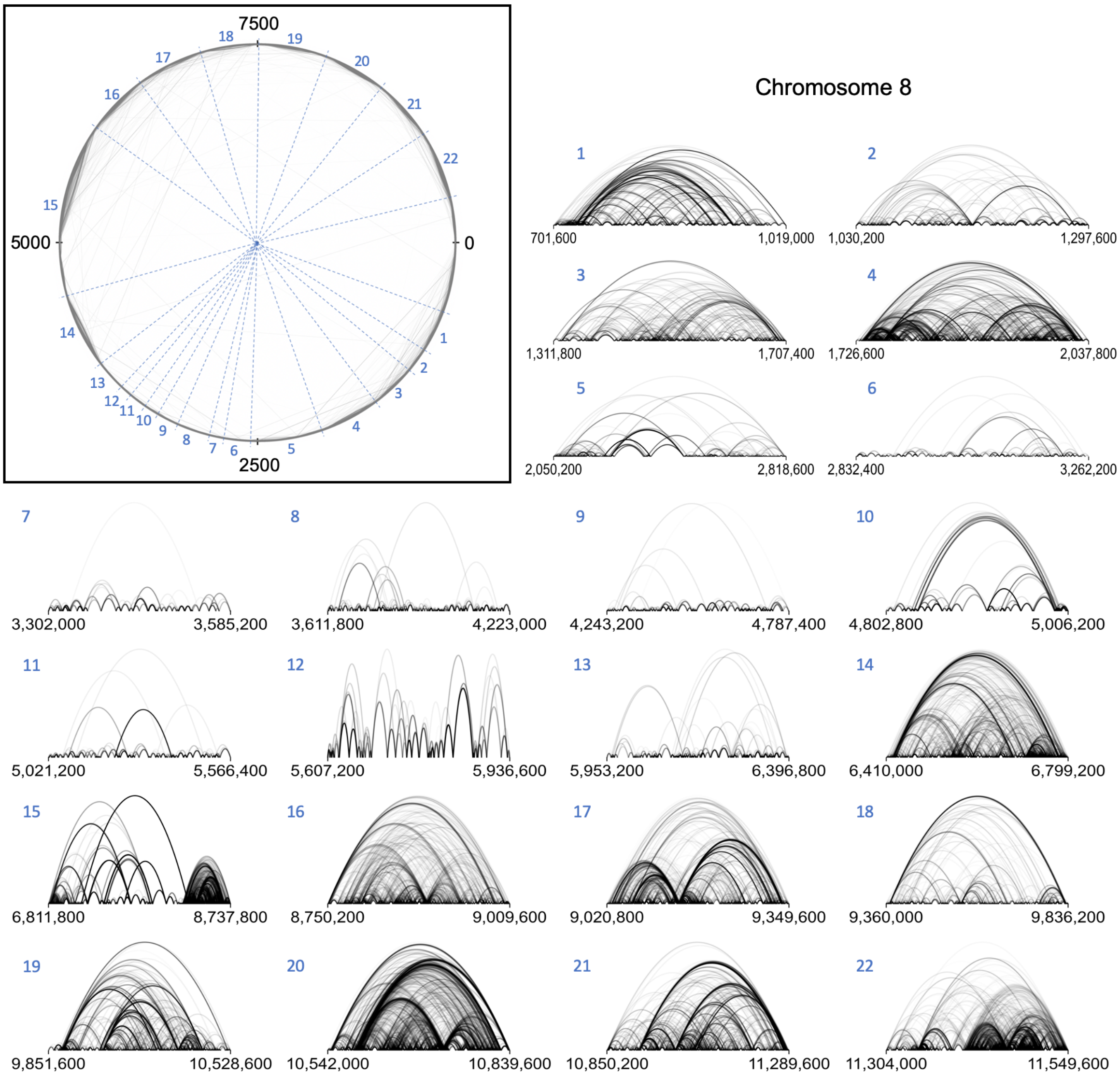


Figure S5: Majority of attribution scores on Micro-C contacts are attribution scores of contacts within topologically associating domains (TADs).

The Mirco-C contacts within the first 10k sequences in Chromosome 8 are visualized in a circle. 0.988 of the total attribution scores for all chromatin features are total attribution scores of contacts within TADs, and 0.982 of the contacts are in TADs. The blue dashed lines show the hESC TAD boundaries. The black numbers on the circle index the 10k sequences, and the blue small numbers index the 22 TADs. The attribution scores of contacts for all chromatin features within each TAD are plotted. The color transparency of the lines represents the values of attribution score.

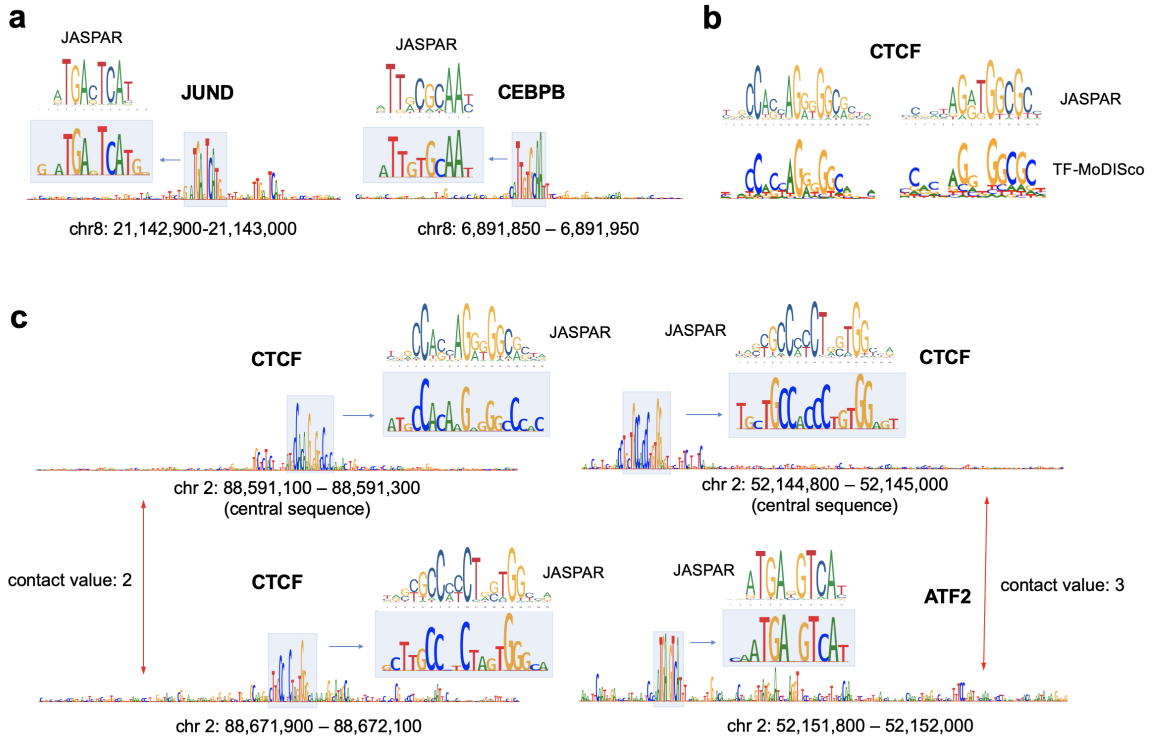


Figure S6: Visualization of attribution scores on DNA sequences.
(**a**) Attribution scores of DNA sequences for two specific TFs, JUND and CEBPB. The height of each letter (A,T,C,G) shows the attribution score for the exact base pair. The high score regions are compared with known motifs from the JASPAR database [22]. (**b**) CTCF motifs generated by TF-MoDISco [23] using attribution scores of 100 sequences whose CTCF bidning sites are successfully predicted by ECHO. Two related patterns are found by TF-MoDISco. (**c**) Visualization of attribution scores of the central sequences and the neighbor sequences toward TFs binding on central sequence. The TF binding events given by peak bed files and the central sequences are marked out. The Micro-C contact values between sequences are given. The correlated high attribution score regions reveal the collaborative binding mechanisms of TFs.

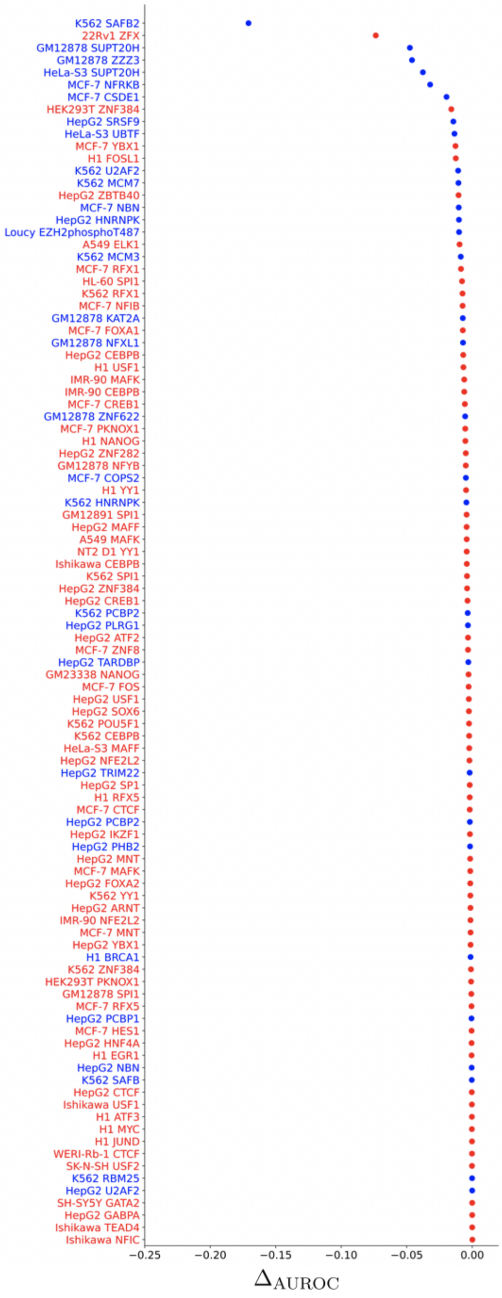


Figure S7: 100 TFs with the cell lines which have lowest performance improvement or perform even worse compared to DeepCNN. TFs without known motifs from JASPAR are marked in blue, others are marked in red.

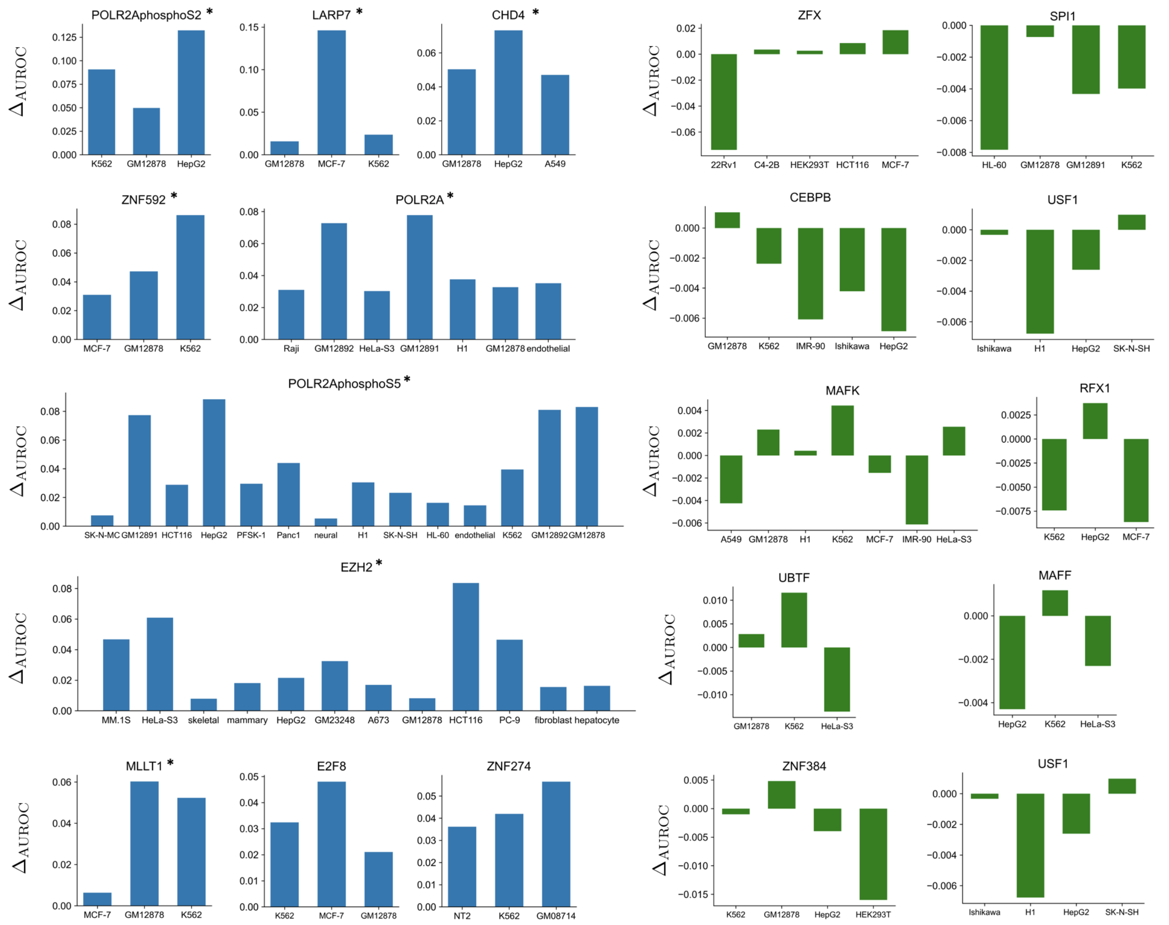


Figure S8: Differences of AUROC scores comparing ECHO with DeepCNN for TFs in multiple cell lines.
We identify ten TFs for which ECHO and DeepCNN predict quite differently among more than three cell lines. The $Y$-axes show the differences of AUROC scores (AUROC from ECHO minus AUROC from DeepCNN). TFs without known motifs are marked with '$*$'. (Left panels) TFs whose AUROC scores are significantly higher in ECHO than DeepCNN. (Right panels) TFs whose AUROC scores are slightly higher or lower in ECHO than DeepCNN.
